# An *in vitro* human-based fracture gap model – Mimicking the crosstalk between bone and immune cells

**DOI:** 10.1101/2020.06.26.165456

**Authors:** Moritz Pfeiffenberger, Alexandra Damerau, Igor Ponomarev, Christian H. Bucher, Yuling Chen, Dirk Barnewitz, Christa Thöne-Reineke, Paula Hoff, Frank Buttgereit, Timo Gaber, Annemarie Lang

**Affiliations:** Charité – Universitätsmedizin Berlin, corporate member of Freie Universität Berlin, Humboldt-Universität zu Berlin, and Berlin Institute of Health, Department of Rheumatology and Clinical Immunology, Berlin, Germany; German Rheumatism Research Centre (DRFZ) Berlin, a Leibniz Institute, Berlin, Germany; Research Center of Medical Technology and Biotechnology, Bad Langensalza, Germany; Charité – Universitätsmedizin Berlin, corporate member of Freie Universität Berlin, Humboldt-Universität zu Berlin, and Berlin Institute of Health, Berlin Institute of Health Center for Regenerative Therapies, Berlin, Germany; Charité – Universitätsmedizin Berlin, corporate member of Freie Universität Berlin, Humboldt-Universität zu Berlin, and Berlin Institute of Health, Julius Wolff Institute, Berlin, Germany; Institute of Animal Welfare, Animal Behavior and Laboratory Animal Science, Department of Veterinary Medicine, Freie Universität Berlin, Berlin, Germany; Endokrinologikum Berlin, MVZ am Gendarmenmarkt, Berlin, Germany

**Keywords:** bone, fracture, fracture hematoma, immune cells, *in vitro* model

## Abstract

The interaction between the bone and immune cells plays a crucial role in bone pathologies such as disturbed fracture healing. After a trauma, the initially formed fracture hematoma in the fracture gap contains all important components (immune/stem cells, mediators) to directly induce bone regeneration and is therefore of great importance but most susceptible to negative influences. Thus, reliable *in vitro* models are needed to study the underlying mechanisms and to predict the efficiency of novel therapeutic approaches. Since common bioengineering approaches exclude the immune component, we introduce an *in vitro* 3D fracture gap model which combines scaffold-free bone-like constructs with a fracture hematoma model consisting of human peripheral blood (immune cells) and bone marrow-derived mesenchymal stromal cells. Our *in vitro* 3D fracture gap model provides all osteogenic cues to induce the initial bone healing processes, which were further promoted by applying the osteoinductive deferoxamine (DFO). Thus, we were able to distinctly mimic processes of the initial fracture phase and demonstrated the importance of including the crosstalk between bone and immune cells.

## Introduction

Bone is an essential part of the musculoskeletal system shaping the body and enabling locomotion and stability. Bone pathologies such as osteoporosis or disturbed fracture healing lead to pain, immobility, inflexibility, considerable loss of life quality and even mental illnesses^1^. Traumatic events can result in bone fracturing accompanied by vessel ruptures and the opening of the bone marrow channel. The pivotal event in the initial phase of fracture healing is the formation of the fracture hematoma, which mainly consists of immune and progenitor cells^2,3^. Negative influences on the initial phase by medications or comorbidities such as diabetes, rheumatoid arthritis or immunosuppression can lead to disturbed fracture healing occurring in approximately 10% of patients with fractures^4,5^. Recent treatment strategies have achieved high technology standards with regard to fixation systems such as plates or implants, regenerative approaches using autologous bone graft (gold-standard) or the additional application of stem cells and/or growth factors^6^. Therefore, preclinical studies are highly needed to tackle the unmet clinical need, especially with respect to an aging population and the increase of comorbidities.

The surgical removal of the fracture hematoma results in a prolonged healing process, while transplantation in an ectopic location elevates bone formation^7^. Formation of the fracture hematoma and the constricted interplay of pro- and anti-inflammatory processes are considered as the starting point of bone regeneration^8-10^. Since the bone marrow cavity is opened during the fracture, the bone marrow acts as a resource for chondro- and osteo-progenitor cells such as mesenchymal stromal cells (MSCs). Therefore, it can be hypothesized that osteogenic induction within the fracture gap is directly induced and controlled by signals from bone components in the vicinity of the fracture gap^11,12^. Hence, the crosstalk between immune cells from peripheral blood (after vessel rupture) and the bone marrow, and osteo-progenitor cells is essential and needs to be considered in preclinical studies^13^. The today’s gold-standard of preclinical drug, compound screening and risk assessment is the use of animal models - mainly rodents (mice and rats) - which is in accordance with most national legal requirements. Nevertheless, trans-species differences may be responsible for the limited transferability of findings to the human patient^14,15^. Mimicking the *in-patient* situation in preclinical studies is highly encouraged and evading cross-species differences by novel *in vitro* approaches is of great interest. During the past decades, conventional *in vitro* cell culture systems have been revised and improved to provide more physiological and human-relevant features. This development was mainly driven by the triumph of regenerative medicine relying on improvements in tissue engineering to produce e.g. huge batches of primary cells, 3D nature-resembling artificial tissues or biocompatible biomaterials. Furthermore, the rapid technical evolution allocating sophisticated biomaterials, bioreactors and microfluidic platforms allows the development of innovative human-relevant *in vitro* systems as alternative or predictive support to animal testing^16^.

Current *in vitro* systems focus on mimicking bone development, endochondral ossification or the bone homoeostasis itself by using spheroids, scaffold-based or scaffold-free model systems. Common cell sources are either primary bone-related cells such as osteoblasts, osteocytes, osteoclasts or MSCs as progenitor cells^16,17^. To mimic fracture healing, models mainly focus on later stages of the regeneration processes particularly endochondral ossification or remodeling. We have previously described the development of a fracture hematoma model consisting of human whole blood and a certain amount of human (h)MSCs^18^. However, to study the initiated processes in an interconnected manner and more adequate experimental setting, the combination of the bone components with the fracture hematoma (immune component) remains elusive.

Within our study, we have developed and characterized scaffold-free bone-like constructs (SFBCs) based on mesenchymal condensation which exceeded the dimensions of spheroids. These SFBCs were co-cultivated with *in vitro* fracture hematoma (FH) models to (i) confirm the capability of the SFBCs to act as an osteogenic inducer, (ii) to mimic the initial phase of fracture healing with adequate culture conditions and (iii) to use the system as a platform to test potential therapeutics (DFO - deferoxamine).

## Results

### SFBCs are characterized by permeating mineralization

MSCs are well-known for their pronounced osteogenic capacity, especially when cultivated *in vitro*^17,19^. However, in a first step, we wanted to know if it is possible to employ mesenchymal condensation as a macroscale approach with consistent 3D self-organization and permeating mineralization. Thus, SFBCs were generated by hMSC condensation and treatment with osteogenic medium until analysis at week 12 (**Fig 1a**). Macroscopic observation indicated comparable generation of SFBCs from different hMSC donors with a diameter of approx. 1 cm and a thickness of 0.5 cm (**Fig. 1b**). To verify the mineralization, *in vitro* computed micro-tomography (µCT) was performed, showing a consistently high mineralization in the outer area which was slightly reduced towards the center (**Fig. 1c**). 3D reconstruction yielded the presence of mineralized tissue as indicated by parameters such as bone volume (BV; mean = 5.1 ± 3.7 mm^3^) and bone surface (BS; mean = 276.1 ± 195.4 mm^2^) (**Fig. 1d**). To quantify the connectedness of the mineralized areas, we additionally examined the trabecular pattern factor (TBPf). This parameter was originally invented to evaluate trabecular bone. Although no osteoclasts were present in the SFBCs, thus the formation of clear trabeculae was not expected, we use this parameter to distinguish between concave (= connected) and convex (= isolated) structures. Low or even negative values represent hereby high connected tissue which was found in at least 4 (TBPf < 2) out of 9 SFBCs (**Fig. 1d**). To evaluate the structural morphology of the SFBCs in further detail, we used scanning electron microscopy (SEM) and found similar morphology when compared to human native bone (**Fig. 1e, f**). In detail, the top view shows strong matrix formation and a closed superficial layer with certain, isolated crystal-like depositions while the cut face revealed a layer-like structure.

**Figure 1:**
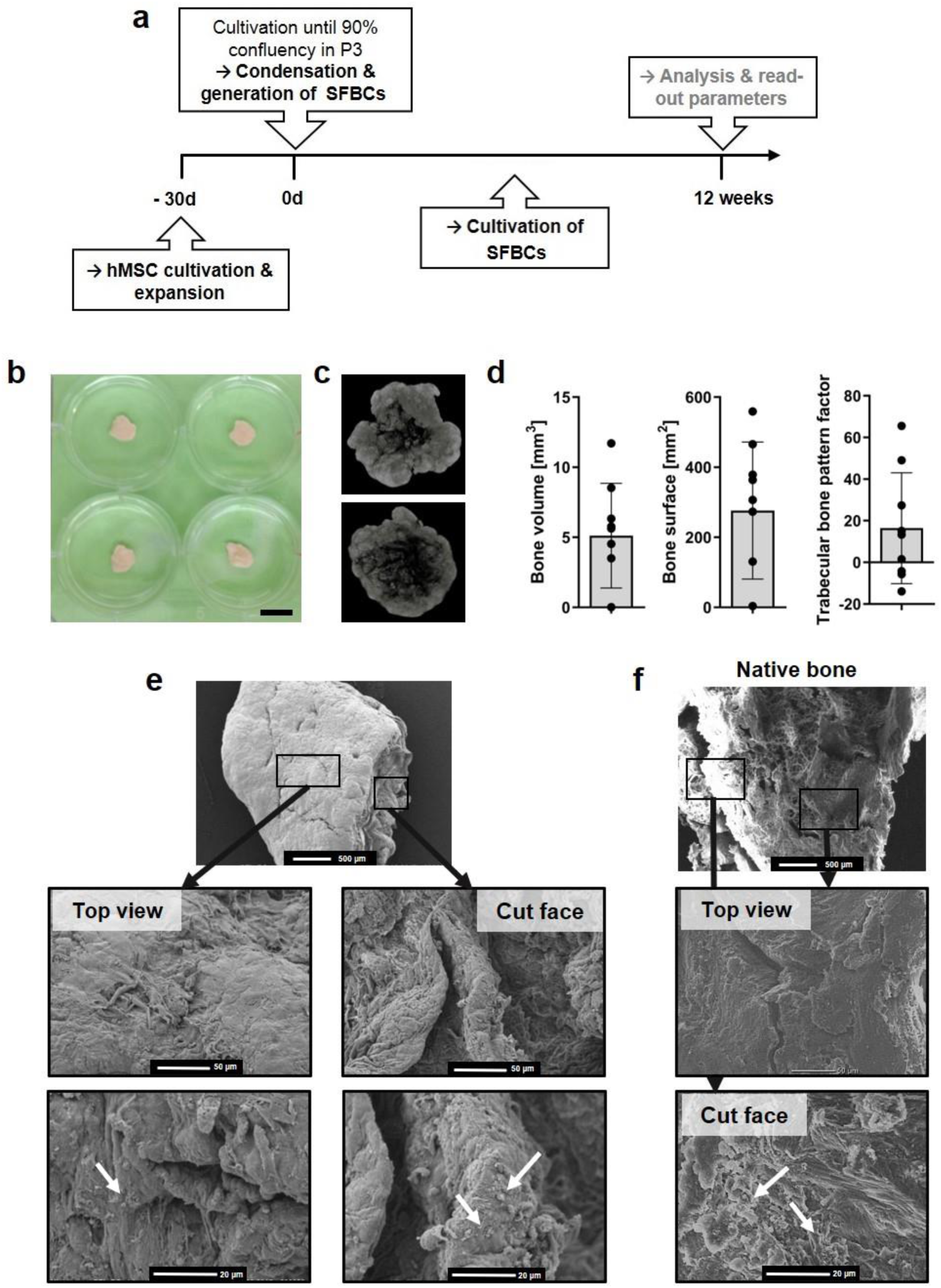
Characterization of the SFBCs with respect to mineralization and structure. (**a**) Study design; (**b**) Exemplary images of SFBCs. Scale bar indicates 1 cm. (**c**) 3D reconstruction of µCT. Exemplary images of n= 9. (**d**) Quantitative results from µCT analysis. Data are shown as mean ± SD. n= 9. (**e**) Structural/morphological examination of the SFBCs in comparison to (**f**) native bone using scanning electron microscopy. Exemplary images of n= 3 SFBCs and n= 2 native human bone pieces. Scale bars are indicated in the images. Arrows mark isolated crystal-like depositions.

To get a more detailed overview on the matrix composition, assorted histological and immunohistochemical stainings were applied. Cells were homogenously distributed within the SFBCs as indicated by hematoxylin and eosin (H&E) staining while Alizarin Red S and von Kossa staining confirmed the deposition of calcium and phosphate, respectively throughout the tissue (**Fig. 2a**). Interestingly, SFBCs which showed a higher amount of negative stained area, showed a more pronounced layer-like structure while the layers itself were strongly mineralized at the borders (**Fig. 2a**; lower row). Negative controls were cultivated without osteogenic medium (**Fig. 2b**). In addition, we observed the expression of *collagen type I* (Col I) and *alkaline phosphatase* (ALP), which are typical markers for osteogenic processes, while no Col II, a typical marker of chondrogenesis, was found (**Fig. 2c**).

**Figure 2:**
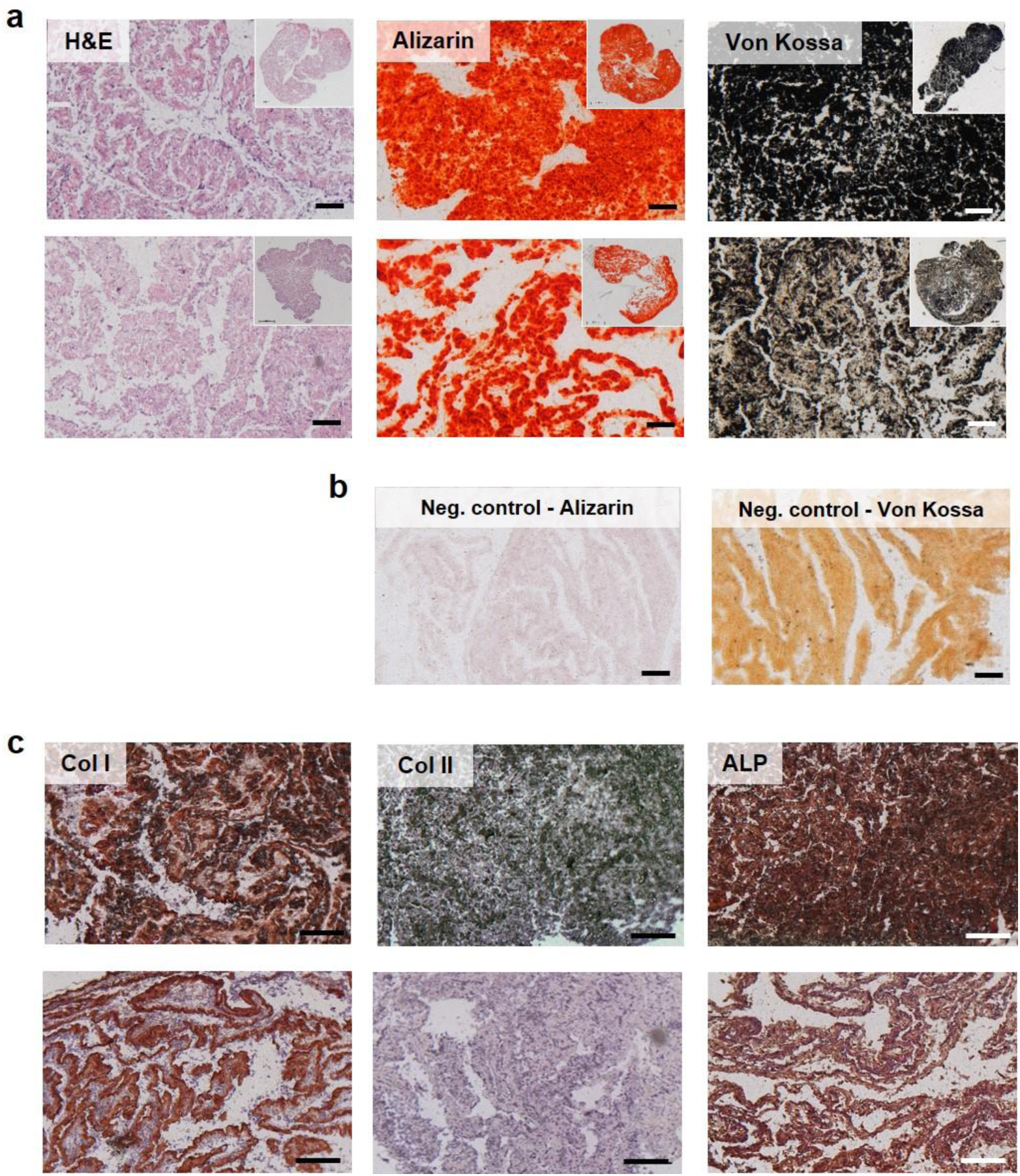
Histological and immunohistochemical examination of the SFBCs. (**a**) Exemplary images of H&E, Alizarin red staining and von Kossa stains. Upper row is exemplary for fully calcified SFBCs and lower row for less fully calcified SFBCs. n= 9. (**b**) Negative control for Alizarin red and von Kossa staining. (**d**) Immunohistochemical stainings for Col I, Col II and ALP. All scale bars indicate 200 µm. All images are exemplary for n=9.

### SFBCs show profound expression of bone-specific markers

In order to confirm the previous findings on a molecular level, immunofluorescence staining for osteopontin (OPN) and osteocalcin (OC), two non-collagenous bone matrix proteins were analyzed and yielded the evenly distributed, distinct protein expression (**Fig. 3a, Fig. 3b**). While OPN is mainly produced by immature osteoblasts, OC is a marker of late stage osteoblasts indicating the presence of different cell states within the SFBCs. mRNA expression analysis showed high levels of osteogenic marker genes such as *secreted phosphoprotein 1* (*SPP1*) and *distal-less homeobox 5* (*DLX5;* **Fig. 3**). *SPP1* was significantly higher expressed (10-fold), while *DLX5* was higher expressed by trend (8-fold) when compared to monolayer MSCs. In contrast the early osteogenic transcription factor *runt-related transcription factor 2* (*RUNX2*) and the chondrogenic marker *SRY-box transcription factor 9* (*SOX9*) were expressed on a more basal level. Interestingly, *receptor activator of NF-κB ligand* (*RANKL)* was highly expressed (**Fig. 3c**). Moreover, *vascular endothelial growth factor* (*VEGFA*) was significantly higher expressed (10-fold), although other hypoxia-inducible factor (HIF1) target genes such as *phosphoglycerate kinase 1* (*PGK1*), *lactate dehydrogenase A (LDHA), endothelial PAS domain-containing protein 1* (*EPAS*) and *HIF1* were comparably or lower (*LDHA*) expressed as in monolayer MSCs (**Fig. 3d**).

**Figure 3:**
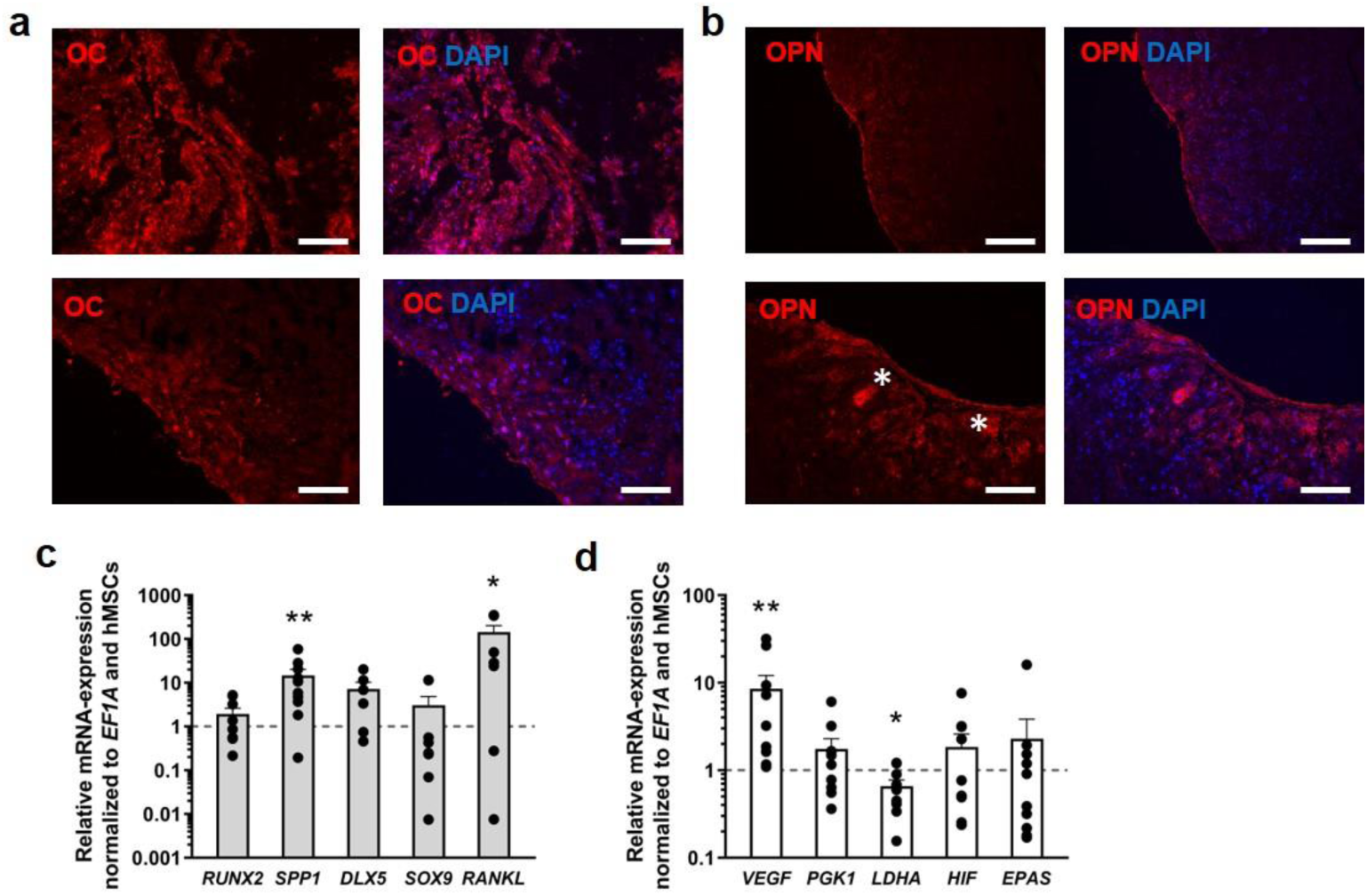
Immunofluorescence and mRNA expression analysis of SFBCs. (**a-b**) Immunofluorescence staining of osteocalcin (OC, **a**) and osteopontin (OPN, **b**). White asterisks highlight cells with high OPN production. To reveal the nuclei of present cells, all slides were counter-stained with DAPI. All scale bars indicate 200 µm. Images are exemplary for n=3. (**c**) qPCR results of mature osteogenic markers (*SPP1, DLX5*) (n=8-10), markers indicative for osteoprogenitors (*RUNX2, SOX9*), and (**d**) metabolic marker *VEGFA* is highly expressed while other metabolic markers (*PGK1, LDHA, HIF1, EPAS*) remain at a basal level (n=10). Data are shown as mean ± SEM. For statistical analysis the results were compared to the expression level of monolayer hMSC and the Wilcoxon signed rank test was used (Table S1); \**p*<0.05, \*\**p*<0.01.

### Co-cultivation indicates biological functionality of SFBCs

To confirm the functional activity of the SFBCs and to mimic the initial phase of fracture healing in an adequate experimental setting, SFBCs were directly co-cultivated with FH models under hypoxic conditions (5% CO_2_, ∼ 1% O_2_) for 48h, since the initial phase of fracture healing in humans takes place within the first 72h^2^ (**Fig. 4a**).

**Figure 4:**
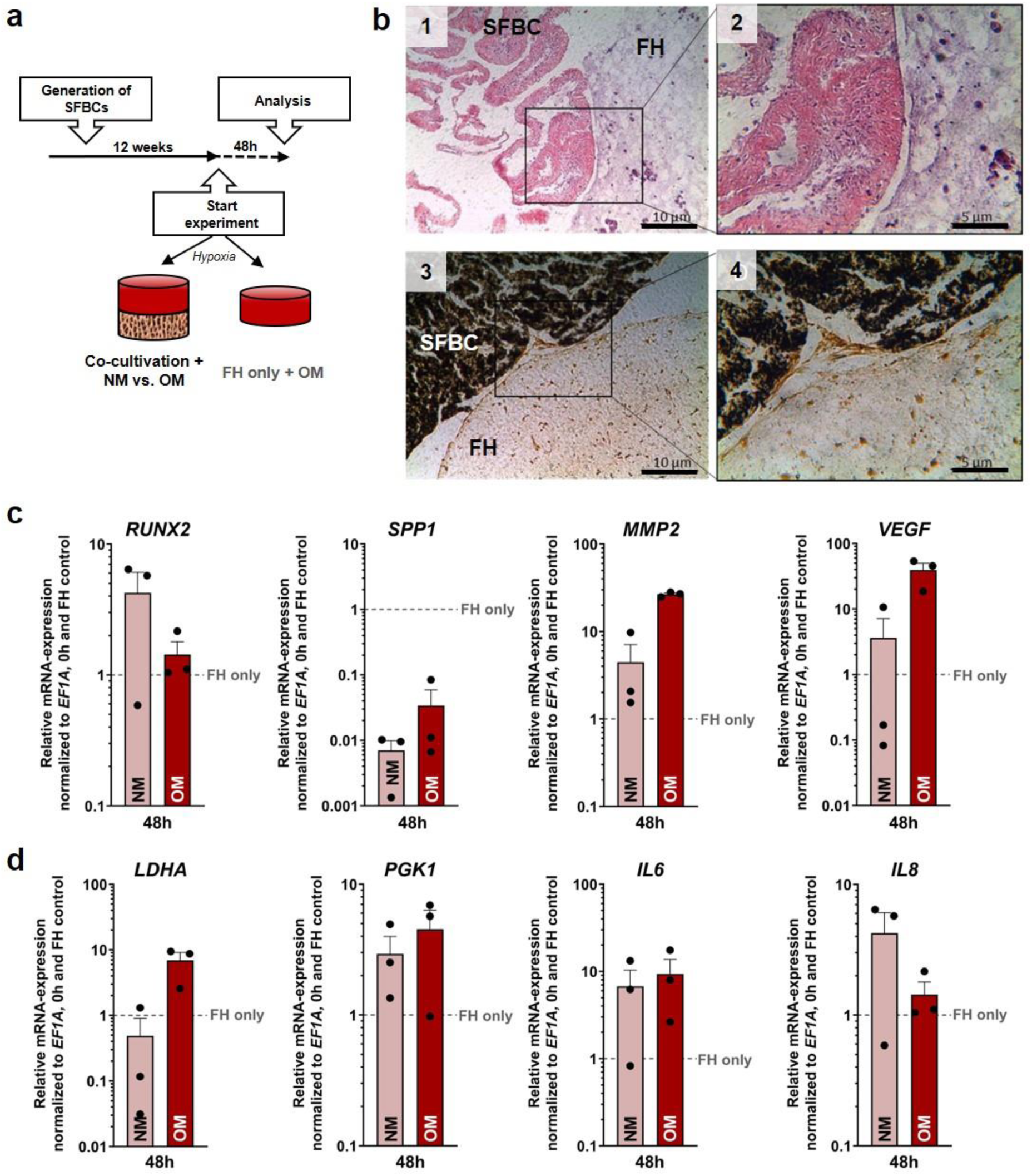
Co-cultivation of SFBCs and *in vitro* FH. (**a**) Experimental setup for the co-cultivation study. (**b**) Exemplary images of H&E (1, 2) and von Kossa staining (3, 4). Images in the right row (2, 4) are magnifications. Scale bars show 10 and 5 µm. (**c, d**) qPCR results of the FH model of the co-cultivation system after 48h co-cultivation under hypoxic conditions. Data is normalized to *EF1A*, 0h and the FH only control group cultured without SFBC in OM under hypoxic conditions (median). Data are presented as mean ± SEM (n= 3). Statistical significance was determined using the Wilcoxon signed rank test (matched-pairs and unpaired to hypothetical value = 1; Table S2 - S5).

Although we have not fixed both models to each other by any technical measures, we observed a close contact of the FH with the corresponding SFBC allowing direct cell-cell-contact and crosstalk between the models. After co-cultivation for 48h, H&E staining revealed the typical cell morphology with no obvious alterations within the SFBCs, while the cells in the FH seemed to be evenly distributed. The calcification throughout the SFBC was reconfirmed via von Kossa staining with no obvious calcification in the border areas of the FH (**Fig. 4b**). We have previously shown that most interesting observations in our *in vitro* FH were visible between 12-48h with respect to immune cell survival and activity^18^. Thus, in a first experiment, mRNA expression in the FH and the SFBC was analyzed after 12 and 48h and normalized to the mean expression at 0h (**Fig. S1 and S2**, respectively). In addition, we compared the cultivation with normal medium (NM) and osteogenic medium (OM, control) to analyze the effect of the additional osteogenic impact on the co-culture system. In short, analysis of the mRNA-expression in the FH indicated a time-consistent expression of almost all genes with only slight differences between NM and OM (**Fig. S1**). The main differences were observed for *matrix metalloproteinase* (*MMP2*), significantly higher expressed at 48h when cultivated with OM. The inflammatory markers *interleukin 6* (*IL6*) and *IL8* were slightly lower expressed after 12h, although gene expression was upregulated in NM and OM after 48h (**Fig. S1**). Within the SFBCs, *SPP1* was elevated after 12h of incubation with NM and was marginally lower expressed in NM and higher expressed in OM at 48h compared to 0h. *MMP2, VEGFA* and *IL8* were highly expressed at both time points with no differences between the cultivation medium. (**Fig. S2**).

Based on this data, further expression analysis and experiments were conducted after 48h of co-cultivation. To verify the osteoinductive potential and biological functionality of the SFBC, we compared the results from the co-cultivated FH model with a FH only control group (treated with OM under hypoxic conditions). The normalization of the data to the starting point (0h) and the FH control revealed a substantial higher expression of *MMP2, VEGF, PGK1* and *IL6* in both groups (NM and OM) when compared to the FH control (**Fig. 4c, d**). While *RUNX2* and *IL8* were higher expressed in the NM group compared to the OM and FH control group *SPP1* was lower expressed in both groups and *LDHA* only in the NM group. Based on these findings, we concluded that the SFBCs show a comparable osteoinductive capacity as the OM medium and biological functionality (**Fig. 4c, d**).

Regarding the protein release, we confirmed the secretion of the pro-inflammatory IL-6 and the pro-inflammatory/-angiogenic IL-8. The pro-inflammatory granulocyte/macrophage stimulating factor (GM-CSF) and the macrophage inflammatory protein MIP were released (**Fig. 5**).

**Figure 5:**
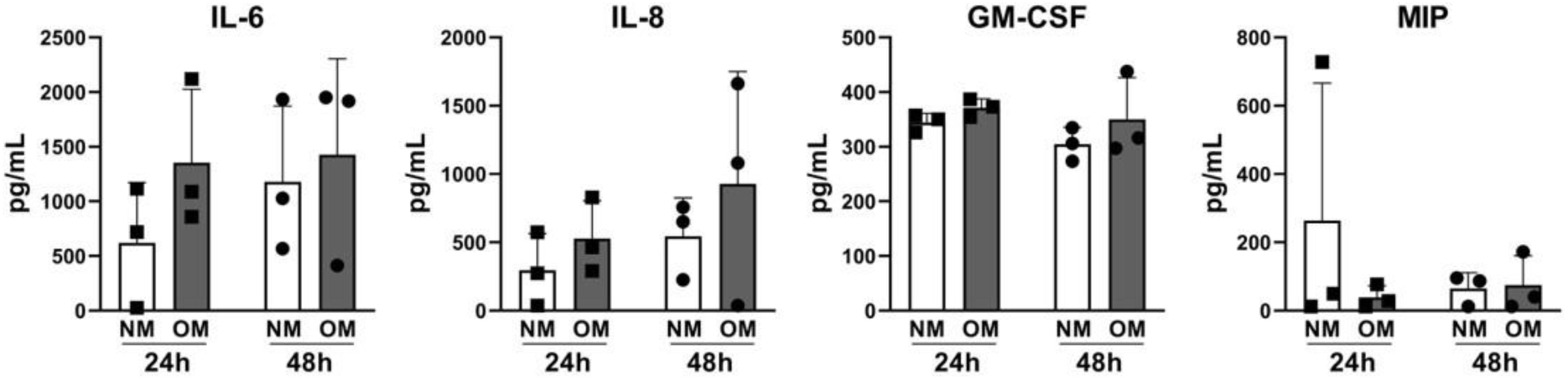
Analysis of the supernatant after co-cultivation. Supernatants were collected after 24h and 48h and analyzed via Multiplex assay. Statistical significance was determined using the Kruskal Wallis test with Dunn’s multiple comparisons test (Table S4). Data is shown as mean ± SD.

### DFO treatment intensifies pro-inflammatory processes

For evaluation of the model suitability as a platform to test potential therapeutics, we supplemented 250 µM DFO for 48h and analyzed the changes in mRNA expression and protein release. DFO is an iron chelator which inhibits prolylhydroxylases to chemically stabilizes HIF and is a well-known osteo-inductive substance^20^. After co-cultivation for 48h in NM, we found that DFO triggered the expression of osteogenic, angiogenic and hypoxia-related genes in the *in vitro* FH with barely any effect on the SFBCs (**Fig. 6a, b**). In detail, the expression of the osteogenic marker *SPP1* in the FH was higher expressed compared to the untreated control, while DFO had barely any effect on the expression of *RUNX2*. The inflammatory markers *MMP2* and *IL6* were additionally elevated while the pro-angiogenic factor *VEGFA* was highly expressed under the influence of DFO compared to the untreated control group, while *LDHA* was also elevated compared to the untreated control (**Fig. 6a**). These results confirm the biological activation capacity of DFO and the possibility to monitor these effects within a short time period. Referring to the mRNA-expression within the SFBCs, we observed a pattern very similar to untreated conditions, which were expected due to the short treatment period (**Fig. 6b**). However, when analyzing the samples treated with OM, we did not see any on the gene expression compared to the untreated control leading to the assumption that the SFBC as natural trigger is more favorable than providing another more artificial (OM) medium (**Fig. 6c, d**). Finally, the protein release of IL-6 and IL-8 was highly induced after 24h NM and DFO treatment (**Fig. 6e**).

**Figure 6:**
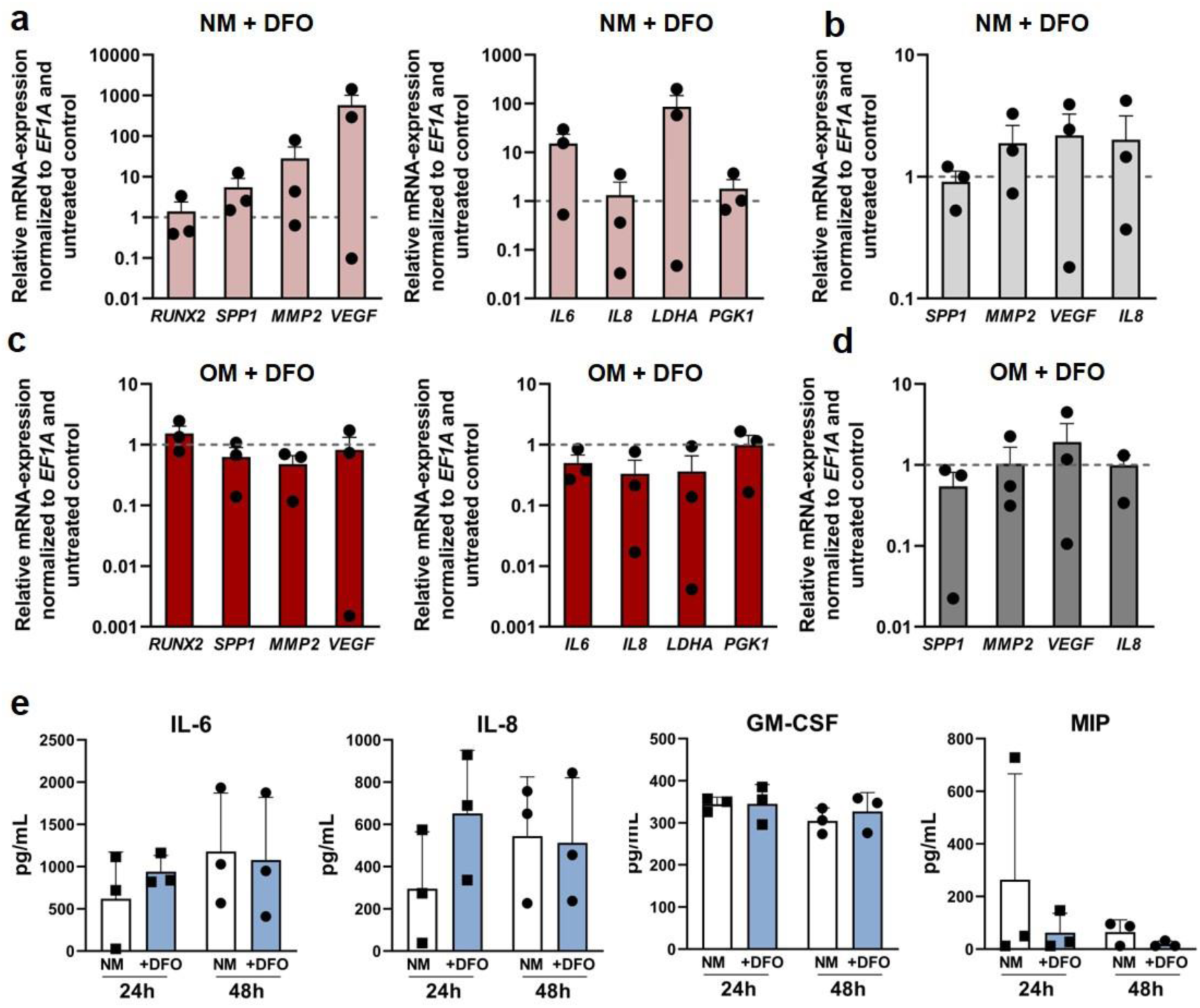
Co-cultivation of SFBCs and *in vitro* FH with supplementation of DFO. (**a**) qPCR results of the FH or the (**b**) SFBCs after 48h co-cultivation with supplementation of NM and 250 µM DFO. Data are normalized to *EF1A* and the untreated control and presented as mean ± SEM (n= 3). Statistical significance was determined using Wilcoxon signed rank test (Table S5, S6). (**c**) qPCR results of the FH or the (**d**) SFBCs after 48h co-cultivation with supplementation of OM and 250 µM DFO. Data are normalized to *EF1A* and the untreated control and presented as mean ± SEM (n= 3). Statistical significance was determined using Wilcoxon signed rank test (Table S7, S8). (**e**) Supernatants were collected after 24h and 48h and analyzed via Multiplex assay. Statistical significance was determined using Kruskal Wallis test with Dunn’s multiple comparisons test (Table S9). Data is shown as mean ± SD.

## Discussion

Our data show that the presented 3D *in vitro* fracture gap model is able to distinctly recapitulate key features of the initial phase of fracture healing. Therefore, we first developed and characterized SFBCs based on mesenchymal condensation, which were subsequently co-cultivated with FH models.

Improved tissue engineering approaches employ mesenchymal condensation as natural form of 3D self-assembly or self-organization consisting exclusively of the cells and their own produced extracellular matrix (ECM)^21^. MSC from the bone marrow, but also the adipose tissue as well as primary cells (osteoblasts) or induced pluripotent stem cells have been used depending on the availability and phenotype stability. It has been described that bone marrow-derived MSCs tend to rather mineralize than undergo chondrogenesis after mesenchymal condensation^17,19^. Several different techniques are exerted such as the use of low attachment plates to induce spontaneous MSC aggregation, membrane-based aggregation (e.g. chitosan) or forced aggregation (via centrifugation)^22^.

The applied technique patented by Ponomarev *et al*.^23^ exploits the capacity of MSC to undergo mesenchymal condensation on a macroscale to produce macro-tissues in a highly reproducible standardized manner without causing necrosis formation in the center^24^. The advantage of such macroscale approaches is the physiologically relevant size, geometry, low cell number and density compared to the matrix and mechanical properties when compared to e.g. spheroids^24^. However, the generation and examination of these constructs are time-consuming and require high numbers of cells limiting the throughput of the system but enabling the performance of several analyses from one model. Here, we reported the bone-like structure and ongoing calcification/mineralization of the SFBCs that were verified by *in vitro* µCt (**Fig. 1**) and histology/immunohistochemistry, showing pronounced expression of ALP and Col I in the absence of Col II (**Fig. 2**). These findings are comparable to other studies using MSC aggregates analyzed after 1 to 3 weeks^25,26^. The morphology and structure of the SFBCs resemble immature woven bone with regard to the body structure. To identify bone-representative cell types, we applied immunofluorescence staining. Since osteoblasts and osteocytes are the characteristic cells in native bone, we stained for OPN as well as OC. OPN - a glycol-phosphoprotein^27^ - is mainly expressed in immature osteoblasts and OC can be found in late stage osteoblasts currently transforming to osteocytes. Thus, we have heterogenic differentiation and maturation states within the SFBCs, indicating a heterogeneous cell population in functional balance. With respect to the mRNA expression, we observed augmented expression of osteogenic relevant markers such as *SPP1, DLX5* and *VEGFA* (**Fig. 3**). *SPP1*, the coding gene for OPN, is hereby the most distinct upregulated gene, coherent with the expression of OPN within the immunofluorescence staining. Muraglie *et al*. observed comparable trends including a two to eightfold increase in OC and OPN expression in MSC aggregates cultivated in low attachment plates^28^. Since *RUNX2* is an early upstream transcription factor, high expression on mRNA-level is expected 3 to 7 days after osteogenic induction^26^. After three weeks, the process of ossification and cellular differentiation towards the osteogenic lineage is already in an advanced state^29,30^. *VEGFA* is an essential coordinator, not only of angiogenetic processes and important for fracture healing, but also in the process of endochondral ossification^31^ and is known to enhance osteogenic differentiation *in vitro*^32^. Interestingly, other HIF1 target genes such as *PGK1, EPAS* and *HIF1* were comparably expressed as in monolayer MSCs, also *LDHA* was downregulated (**Fig. 3e**). Since the SFBCs were not cultivated under hypoxic conditions, the increased expression of *VEGFA* might result from an alternative pathway, e.g. induced by transforming growth factor beta 1 (TGF-β1)^33^. *DLX5*, an important transcription factor in osteogenesis and bone development^34^, is also highly expressed within the SFBCs, also indicating an intense ossification process. RANKL is also expressed in mature osteoblasts, differentiating into osteocytes and also regulates osteoblastogenesis indicating the presence of late-stage osteoblasts^35^.

After characterizing their bone-like quantities, we co-cultivated the SFBCs with *in vitro* FHs in order to evaluate the capability of the SFBCs to act as an osteogenic inducer and to recapitulate key features of the initial phase of fracture healing closely to the *in vivo* situation. Previously, we developed an *in vitro* FH model, incubated in osteogenic induction medium, which closely reflects the *in vivo* situation^18,36^. One of the main findings was the importance of hypoxia. Therefore, we included hypoxic conditions in our co-cultivation setup. The co-cultivation of the *in vitro* FH and SFBCs in NM for up to 48h under hypoxic conditions revealed significant initiation of ongoing cellular processes (mRNA-level) for adaptation to hypoxia and osteogenic induction within the FH (**Fig. S1, S2**). These findings are in accordance with results from an *ex vivo* study and an *in vitro* FH model conducted in our group (**Table 1**)^18,37^.

**Table 1:** Gene expression data from an *ex vivo* study using primary human fracture hematomas obtained between 48 and 72 h after trauma (n=40), the i*n vitro* human FH model (n=12, 48 h incubation under hypoxia and the FHs of the *in vitro* fracture gap model (n=3, 48 h incubation under hypoxia in normal medium).

| Gene symbol | *ex vivo* FHs<br>(< 72 h) | *in vitro* human FHs<br>(48 h hypoxia) | *in vitro* fracture gap<br>model (48 h hypoxia) |
| --- | --- | --- | --- |
| *RUNX2* | ↑ | ↑** | ↑ |
| *SPP1* | ↑* | ↑**** | ↑ |
| *VEGFA* | ↑* | ↑*** | ↑ |
| *IL8* | ↑*** | ↑** | ↑ |
| *IL6* | ↑*** | ↑ | ↑ |
| *LDHA* | ↑** | ↑*** | ↑ |
| *MMP2* | n.a. | ↑* | ↑ |

Based on the comparative analysis with an FH control, we concluded that the SFBCs show a comparable osteoinductive capacity as the OM medium and biological functionality indicated by e.g. *VEGF* and *MMP2* (**Fig. 4**). Interestingly, *SPP1* was noticeably lower expressed in the FH when co-cultivated with SFBCs compared to the FH control, which could be explained by either the higher amount of OPN provided by the SFBC (**Fig. 3**) or the considerable high expression/concentration of IL-6 (**Fig. 4 and 5**), as previously reported^38^. The level of pro-inflammatory cytokines (e.g. IL-6 and IL-8) was abundant, which has been also observed in *ex vivo* samples from patients (**Fig. 5**)^2,5,37^. The microenvironment of the fracture hematoma is described by hypoxia, high lactate and low pH due to the disruption of vessels, cell death of e.g. erythrocytes and the lack of nutrients. This cytotoxic environment needs to be counter-regulated to allow the invasion of regenerative cells. Therefore, we included whole human blood instead of isolated peripheral blood mononuclear cells (PBMCs). At the initial stage, the microenvironment in the fracture hematoma is acidotic and switched to neutral and slightly alkaline during the regeneration process^39^ which can be triggered by hypoxia. Interestingly, upon co-cultivation *LDHA* was highly upregulated in the FH model (**Fig. S1**).

Furthermore, the effect of certain immune cells during the initial phase of fracture healing has been studied in detail during the last decade. Lymphocytes play a crucial role as shown in RAG1(-/-) mice supposing a detrimental effect of adaptive immune cells^40^. A negative impact was reported for the presence of CD8^+^ cytotoxic T-cells in humans and mice^41^, while CD4^+^ cells have been shown to enhance osteogenic differentiation *in vitro* and upregulated osteogenic markers e.g. *RUNX2* or *OC*^42^. This clearly indicates the need to combine bone and immune cells in *in vitro* approaches to recapitulate the crosstalk, environment and key features of the *in vivo* situation.

With respect to the DFO treatment, we found an upregulation of *HIF*-target genes (*VEGFA* and *LDHA*) indicating the effectivity of DFO to stabilize *HIF* (**Fig. 6**). In addition, pro-inflammatory processes were more pronounced, which is in accordance with current findings in a mouse-osteotomy-model revealing the activation of e.g. *C-X-C motif chemokine ligand* 3 (*Cxcl3*) or *metallothionein 3* (*Mt3*) expression at day 3 after application of DFO in the fracture gap^20^. We did not expect a strong upregulation of osteogenic markers or changes in the SFBCs, which normally require longer treatment periods^29,43^.

Nevertheless, there are many ways to foster the current approach. In order to improve the current protocol to generate SFBC, different approaches such as the addition of specific growth factors, e.g. BMPs, fibroblast growth factors (FGFs) or VEGFs, enhancing the differentiation and bone formation *in vitro* can be considered to fasten up the generation period in the future. Furthermore, bioreactors can be used to approximate the environment to the actual *in vivo* situation by dynamic culturing and restrained environment, while overcoming the lack of nutrient transfer and combining cells with scaffolds^44^. Bioreactors also provide the possibility to withdraw toxic and cell apoptotic signals perhaps allowing a longer cultivation period of the fracture gap model.

Taken together, within our 3D *in vitro* fracture gap model, we have been able to distinctly mimic key features of the initial phase of fracture healing which can be used i) to study potential underlying mechanism of fracture healing disorders, especially with respect to immunologically restricted patients^4,5^, requiring the crosstalk between immune cells and bone and ii) as a prediction tool for potential new therapeutic strategies actively implementing the 3R principle.

## Material & Methods

### Bone marrow derived MSC isolation, cultivation and characterization

Human mesenchymal stromal cells (hMSC) were isolated from bone marrow of patients undergoing total hip replacement (registered and distributed by the “Tissue Harvesting” Core Facility of the Berlin Institute of Health Center for Regenerative Therapies (BCRT); donor list in **Table 2**). All protocols were approved by the Charité-Universitätsmedizin Ethics Committee and performed according to the Helsinki Declaration (ethical approval EA1/012/13). MSC isolation was performed as described in detail before^18,45^. Briefly, bone marrow was transferred cell culture flask, covered with normal expansion medium (NM) containing DMEM+GlutaMAX (Gibco), 10 (v/v) % FCS (Biowest), 1 (v/v) % Penicillin/Streptomycin (Gibco), 20 (v/v) % StemMACS MSC Expansion Media XF (Miltenyi Biotech) and incubated at 37°C in 5% CO_2_ atmosphere (app. 18% O2). Medium was changed after 3-4 days when cells became adherent. Isolated cells were expanded in at 37 °C, 5% CO_2_. Medium exchange was performed weekly and passaging with Trypsin-EDTA (Gibco) was conducted at a cellular confluency of 80-90%. For characterization, MSCs were evaluated at passage 3 for their differentiation potential (osteogenic, adipogenic) and the presence and absence of specific cell surface markers (MSC Phenotyping Kit, Miltenyi Biotech) as described in detail before^18,45^. Human EDTA-blood was collected from healthy donors with written consent^18^.

**Table 2:** hMSC and blood donor information and conducted experiments.

| Donor | Age | Sex | Type of experiments | Used methods |
| --- | --- | --- | --- | --- |
| MSC 1 | 71 | m | Characterization of SFBCs | μCt, gene expression analysis, histology, immunofluorescence |
| MSC 2 | 77 | m |  |  |
| MSC 3 | 69 | m |  |  |
| MSC 4 | 69 | w |  | μCt, gene expression analysis, histology |
| MSC 5 | 81 | w |  |  |
| MSC 6 | 49 | w |  |  |
| MSC 7 | 59 | m |  |  |
| MSC 8 | 80 | w |  |  |
| MSC 9 | 51 | m |  |  |
| MSC 10 | 75 | m | Characterization of SFBCs, Co-cultivation experiments | μCt, gene expression analysis, histology, gene expression analysis, DFO treatment studies |
| MSC 11 | 68 | w | Co-cultivation experiments | Gene expression analysis, DFO treatment studies |
| MSC 12 | 56 | m |  |  |
| Blood 1 | 37 | M |  |  |
| Blood 2 | 26 | M |  |  |
| Blood 3 | 38 | M |  |  |

### Fabrication of the 3D bone-like scaffold-free constructs (SFBCs)

SFBCs were produced based on a patented protocol (patent no.: EP1550716B1) ^46^ which was modified by applying osteogenic medium to induce osteogenic differentiation after 1 week and no application of biomechanical loading to avoid matrix destruction due to mineral formation. The osteogenic medium for generation and maturation contained DMEM/F-12, 10% (v/v) FCS, 1% (v/v) streptomycin/penicillin, 10 mM β-glycerophosphate and 10 nM dexamethasone (Sigma Aldrich). Approx. 10-20 × 10^5^ hMSC/cm^2^ were cultivated in expansion medium until reaching confluency and forming a monolayer cell sheet. These cell sheets were detached and centrifuged at 350 *g* for 15 min at RT. Afterwards, the resulting cell aggregates were cultivated for up to one week with medium exchange every day. Cell aggregates were then transferred to a multi-well plate and cultured for up 12 weeks until performing characterization or co-cultivation with the FH model.

### Generation of FH and co-cultivation – The fracture gap model

The FH models were generated as described previously ^18^. In brief, per FH 2.5 × 10^5^ hMSCs per well were centrifuged for 3 min at 300 *g* in a 96-well plate (U-bottom). Afterwards, the cell pellet was resuspended in 100 µL of EDTA-blood and subsequently mixed with a 10 mM CaCl_2_ solution (solved in PBS). After an incubation time of 30 min at 37 °C, 5% CO_2_ the FH were placed on a SFBC with direct contact and transferred into either NM or osteogenic medium (OM) containing NM supplemented with 10 mM β-glycerophosphate and 0.1 mM L-ascorbic acid-2-phosphate.

For the treatment study, 250 µM DFO (Sigma Aldrich) was supplemented to the medium. The generated fracture gap models were incubated under hypoxic (37 °C, 5% CO_2_ and ∼ 1% O_2_ - flushed with N_2_) conditions in a humidified atmosphere for up to 48h.

### In vitro micro computed tomography (µCT)

SFBCs were scanned at a nominal resolution of 8 μm, with a SkyScan 1172 high-resolution microCT (Bruker). X-ray tube voltage was set at 80 kV, 124 µA with maximized power of 10W and a 0.5 mm aluminum filter was employed to reduce beam hardening effects. The scan orbit was 360 degrees with a rotation step of 0.2 degree. For reconstruction the Bruker NRecon software accelerated by GPU was used and Gaussian smoothing, ring artifact reduction, misalignment compensation, and beam hardening correction were applied. XY alignment was corrected with a reference scan to determine the thermal shift during scan time. The CTAn software (Bruker) was used to analyze the total VOI of SFBCs. The threshold for bony tissue was set globally (determined by the Otsu method) and kept constant for all SFBCs. For analysis we used the bone volume (BV), the bone surface (BS) and the trabecular pattern factor (TBPf)^47^ as measured and calculated by the software.

### Scanning electron microscopy (SEM)

Samples were fixated with 2.5 (v/v) % glutaraldehyde (fixation for 10 min), dehydrated with increasing alcohol concentration 30 (v/v) % - 100 (v/v) % (5 steps) and incubation with 100% hexamethyldisilazane (all Sigma Aldrich). Subsequently, the samples were transferred to a sample holder and gold coating was performed with a fine gold coater JFC-1200 (JEOL). For electron microscopy, the JCM-6000 Plus NeoScope (JEOL) was used for imaging, high vacuum was adjusted.

### Histological stainings

The embedding and slice preparation of the SFBCs was conducted according to the Kawamoto *et al*. ^48^ method to prepare slices of undecalcified bones. In detail, the SFBCs were fixated for 6h in a 4 (v/v) % paraformaldehyde solution (PFA; Carl Roth) followed by an ascending sucrose solution treatment (10 (w/v) %, 20 (w/v) % and 30 (w/v) %) for 24h, respectively and afterwards cryo-embedded with SCEM medium (Sectionlab). Slices were produced with a cryotom using cryofilms (Sectionlab) and afterwards air dried for 20 min and fixated with 4 (v/v) % PFA prior to every histological or immunhistological staining on a microscope slide.

H&E staining was performed as described previously^24^. In short, slices were fixed with 4 (v/v) % PFA (10 min), washed with distilled water and stained with Harris’s hematoxylin solution (Merck Millipore). Staining was followed by several washing steps, a differentiation step (0.25 (v/v) % concentrated HCl) and a second staining step in 0.2 (w/v) % eosin (Chroma Waldeck). Staining was finished by differentiation in 96 (v/v) % and 100 (v/v) % ethanol, fixation with xylol and covering with Vitro-Clud (R. Langenbrinck GmbH).

Alizarin Red S staining was conducted by applying slices after fixation and washing to the 2 (w/v) % Alizarin Red S staining solution (Sigma Aldrich; pH = 4.1-4.3) for 10 min. Afterwards slices were washed in distilled water and differentiation was performed in 0.1 (v/v) % HCL solved in ethanol and fixed by washing two times with 100% ethanol before xylol fixation and covering.

Von Kossa staining was conducted according to the following protocol: air drying, fixation and washing as described above, 3% (w/v) silver nitrate solution (10 min), washing step with distilled water, sodium carbonate formaldehyde solution (2 min), washing step with tap water, 5% (w/v) sodium thiosulphate solution (5 min), washing step with tap water and distilled water, ascending ethanol series (70 (v/v) % - 100 (v/v) %), fixation in xylol and covering.

For Col I, II and ALP staining, slices were rehydrated with phosphate buffered saline (PBS) treated with 3 (v/v) % H_2_O_2_ (30 min), washed with PBS, blocked with 5% normal horse or goat serum (Vector Laboratories) in 2 (w/v) % bovine serum albumin (BSA), and incubated overnight with primary antibodies at 4 °C (Col I antibody: ab6308, 1:500, Abcam; Col II antibody, 1:10, Quartett Immunodiagnostika; ALP antibody, ab95462, Abcam). Afterwards, slices were washed and treated with 2 (v/v) % secondary antibody (biotinylated horse anti-mouse IgG antibody – Col I and Col II; biotinylated goat anti-rabbit IgG antibody - ALP, Vector Laboratories) diluted in 2 (v/v) % normal horse/goat serum/2 (v/v) % BSA/PBS (30 min), washed with PBS, incubated with avidin-biotin complex (Vectastain Elite ABC HRP Kit, Vector Laboratories) (50 min), washed with PBS, incubated with DAB under microscopic control with time measurement (DAB peroxidase (HRP) Substrate Kit, Vector Laboratories) and stopped with PBS. For counterstaining slices were washed with distilled water and stained with Mayer’s hematoxylin (Sigma Aldrich), washed with tap water and covered with Aquatex (Merck Millipore). Pictures were taken with the Axioskop 40 optical microscope (Zeiss) using the corresponding AxioVision microscopy software. Von Kossa staining was quantified using ImageJ and the threshold tool to mark positive (black) and negative (brown) stained areas.

### Immunofluorescence

For the immunofluorescence staining, the slides were rehydrated with PBS and blocked with PBS with 5 (v/v) % FCS for 30 min at RT. Primary osteopontin antibody (mouse anti-human, Abcam) or osteocalcin (rabbit anti-human, Abcam) was diluted 1:50 in PBS/5 (v/v) % FCS/0.1 (v/v) % Tween 20 and incubated for 2h. After washing with PBS/0.1% Tween 20, the secondary antibody (donkey anti-goat A568; Life Technologies/Thermo Scientific) was diluted 1:500 in PBS/5 (v/v) % FCS/0.1 (v/v) % Tween 20 and applied for 1h. Pictures were taken with a Keyence fluorescence microscope BZ 9000 (Keyence) using the DAPI, TexasRed and Cy5 channels.

### RNA isolation and quantitative PCR (qPCR)

SFBCs were transferred to RLT-buffer (Qiagen, Germany) with 1% 2-Mercaptoethanol (Serva) and disrupted using the Qiagen Tissue Ruptor (Qiagen). Total RNA was extracted using the RNeasy Fibrous Tissue Mini Kit (Qiagen) according to the manufacturers’ instructions and the RNA concentration was determined using the Nanodrop ND-1000 (Peqlab). RNA was stored at −80 °C until further processing. The same RNA isolation method was preceded for the SFBCs after co-cultivation.

After co-cultivation, the cells of the FH were filtered through a cell strainer (Corning). After centrifugation for 10 min at 300 *g*, the cell pellet was resuspended in 350 µL RLT buffer with 3.5 µL 2-Mercaptoethanol and total RNA was extracted using the Rneasy Fibrous Tissue Mini Kit according to the manufacturers’ instructions.

The cDNA for both the FH and the SFBCs was synthesized by reverse transcription using TaqMan Reverse Transcription Reagents (Applied Biosystems). qPCR was performed using the DyNAmo Flash SYBR Green qPCR Kit (Thermo Fisher) and the Stratagene Mx3000P (Agilent Technologies). Initial denaturation was for 7 min at 98 °C. Afterwards 50 cycles with 5 sec at 98 °C, 7 sec at 56 °C and 9 sec at 72 °C were performed. The melting curve was analyzed through stepwise increasing the temperature from 50 °C to 98 °C every 30 sec. All primers were purchased from TIB Molbiol (**Table S10**). For the gene expression of the SFBCs, data were normalized to the expression of *eukaryotic translation elongation factor 1 alpha 1 (EF1A)* and to the corresponding MSC culture in 2D, using the delta-delta-Ct-method. For the gene expression in the fracture gap model, data were normalized to the expression of *EF1A* using the deltaCt method and 0h using the deltadeltaCt method.

### Cytokine and chemokine quantification in supernatants

Supernatants were immediately stored at −80 °C after 48h co-cultivation. The concentration [pg/mL] of cytokines and chemokines was determined using multiplex suspension assay (Bio-Rad Laboratories) following the manufacturers’ instructions. Following cytokines and chemokines (lower detection limit) were measured: IL-1β (7.55 pg/mL), IL-2 (18.99 pg/mL), IL-4 (4.13 pg/mL), IL-5 (20.29 pg/mL), IL-6 (25.94 pg/mL), IL-7 (16.05 pg/mL), IL-8 (37.9 pg/mL), IL-10 (37.9 pg/mL), IL-13 (7.21 pg/mL), IL-17 (24.44 pg/mL), interferon-gamma (IFNγ, 56.32 pg/mL), tumor necrosis factor-alpha (TNFα, 59.53 pg/mL), monocyte chemotactic protein-1 (MCP-1, 27.02 pg/mL), macrophage inflammatory protein MIP-1β (6.27 pg/mL), granulocyte colony-stimulating factor (G-CSF, 50.98 pg/mL) and granulocyte-macrophage colony-stimulating factor (GM-CSF, 11.82 pg/mL)

### Statistical analysis

Statistical tests were performed using GraphPad Prism Software version 8. Statistical differences towards a hypothetical value were determined by Wilcoxon signed rank test (unpaired). With respect to the co-cultivation studies, differences between two groups were determined with Wilcoxon matched-pairs signed rank test or between more groups with the Kruskal Wallis test with Dunn’s multiple comparisons test. Probability values of *p*<0.05 were considered to be statistically significant (\*\*\**p*<0.001, \*\**p*<0.01, \**p*<0.05). Details on the statistics per Figure are displayed in the Supplementary Information.

## Acknowledgments

The authors would like to thank Manuela Jakstadt for excellent technical assistance. Bone-marrow was provided from the “Tissue Harvesting” Core Facility of the Berlin Institute of Health Center for Regenerative Therapies (BCRT). FACS analyses were performed together with the Core Facility at the German Rheumatism Research Centre. AL, FB, AD, MP and TG are members of Berlin-Brandenburg research platform BB3R and Charité 3^R^. This study was funded by the German Federal Ministry for Education and Research (BMBF) (project no. 031A334). AL is currently being supported by the Joachim Herz Foundation (Add-on Fellowship 2019). The work of TG was funded by the Deutsche Forschungsgemeinschaft (353142848). Funding bodies did not have any role in designing the study, in collecting, analyzing and interpreting the data, in writing this manuscript, and in deciding to submit it for publication.

## Contributions

Study design: AL, TG, MP; Data collection and analysis: MP, AL, IP, AD, CB, YC; Data discussion and interpretation: FB, CTR, AL, TG, MP; Drafting manuscript: MP, AL, TG; Revising manuscript: FB, IP, CTR, PH.

## Supplementary Information

**Table S1:** Results from Wilcoxon signed rank test for Figure 3c-e

| Specification | Wilcoxon signed rank test |  |
| --- | --- | --- |
|  | Hypothetical median = 1<br>*p*-value | 95% CI |
| *RUNX2* | 0.7344 | -0.50 – 68.05 |
| *SPP1* | **0.0039\*\*** | 0.82 – 27.34 |
| *DLX5* | 0.1563 | -0.54 – 19.25 |
| *SOX9* | 0.7422 | -0.99 - 10.51 |
| *RANKL* | **0.0391\*** | -0.99 – 351.10 |
| *VEGF* | **0.002\*\*** | 0.17 – 25.72 |
| *PGK1* | 0.2754 | -0.45 – 2.22 |
| *LDHA* | **0.0195\*** | -0.66 – 0.20 |
| *HIF* | 0.5566 | -0.76 – 2.18 |
| *EPAS* | 0.7695 | -0.82 – 0.03 |

**Figure S1:**
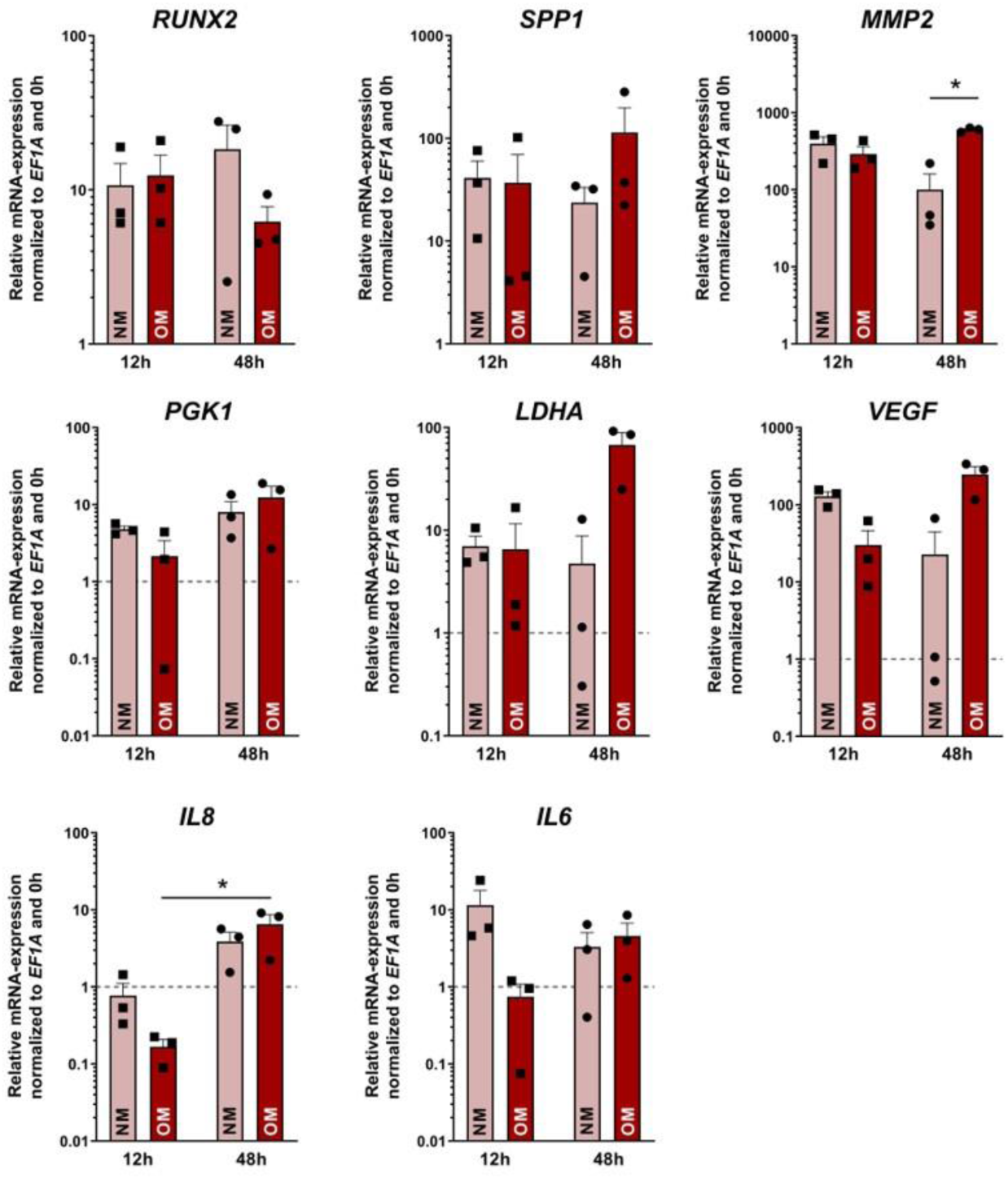
Co-cultivation of SFBCs and *in vitro* fracture hematomas. qPCR results of the fracture hematoma model or the after 12 and 48h co-cultivation under hypoxic conditions. Data are normalized to *EF1A* and 0h and presented as mean ± SEM (n= 3). Statistical significance was determined using the Kruskal Wallis test with Dunn’s multiple comparisons test and Wilcoxon signed rank test (Table S2 - S5). \**p*<0.05.

**Figure S2:**
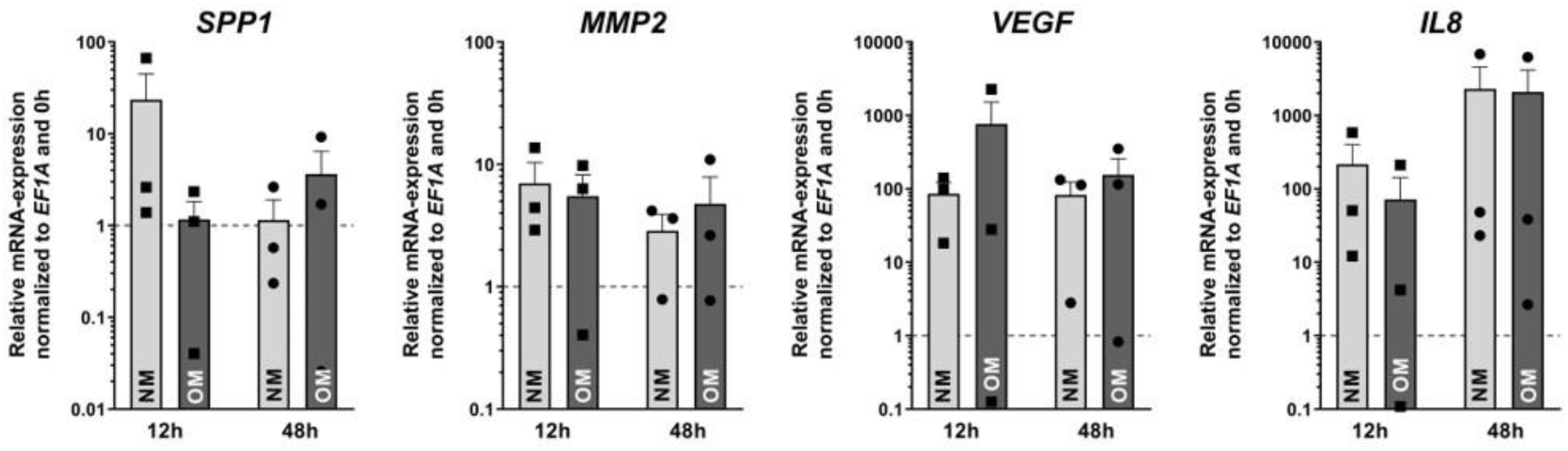
Co-cultivation of SFBCs and *in vitro* fracture hematomas. qPCR results of the SFBCs after 12 and 48h co-cultivation under hypoxic conditions. Data are normalized to *EF1A* and 0h and presented as mean ± SEM (n= 3). Statistical significance was determined using the Kruskal Wallis test with Dunn’s multiple comparisons test and Wilcoxon signed rank test (Table S2 - S5). \**p*<0.05.

**Table S2:**
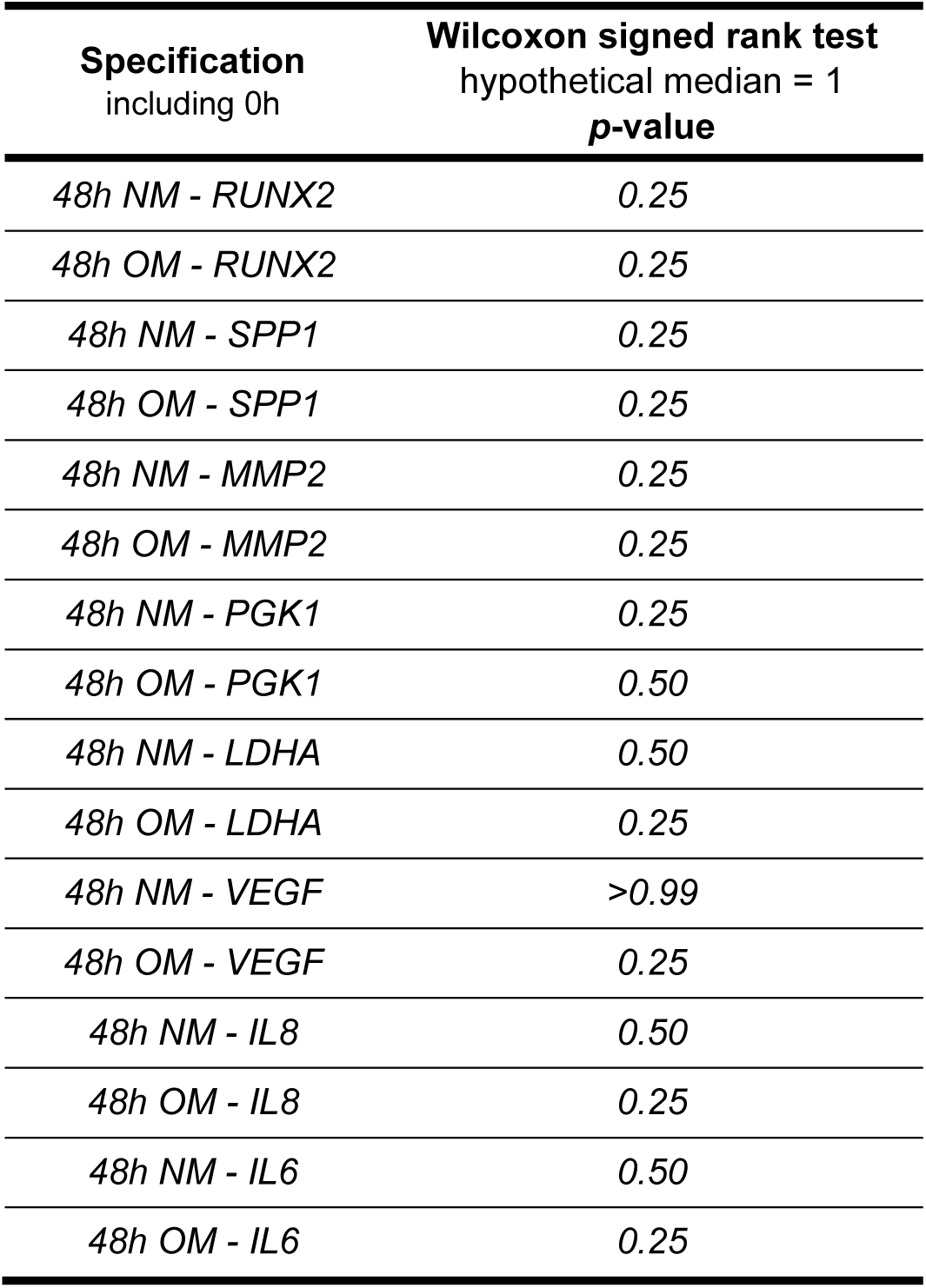
Results from Wilcoxon signed rank test for Figure 4c,d

**Table S3:** Results from Wilcoxon signed rank test for Figure 4c,d

| Specification<br>including 0h | Wilcoxon matched-pairs<br>signed rank test<br>*p*-value |
| --- | --- |
| *NM vs. OM - RUNX2* | *0.50* |
| *NM vs. OM - SPP1* | *0.50* |
| *NM vs. OM - MMP2* | *0.25* |
| *NM vs. OM - PGK1* | *0.50* |
| *NM vs. OM - LDHA* | *0.25* |
| *NM vs. OM - VEGF* | *0.25* |
| *NM vs. OM - IL8* | *0.50* |
| *NM vs. OM - IL6* | *0.75* |

**Table S4:**
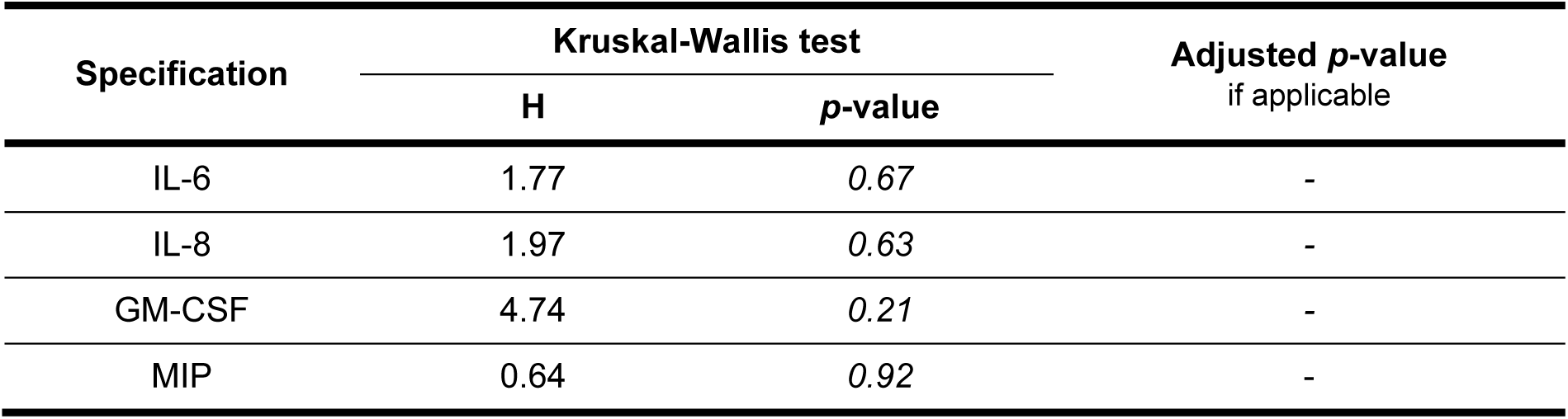
Results from Kruskal Wallis test with Dunn’s multiple comparisons test for Figure 5

| Specification | Kruskal-Wallis test |  | Adjusted *p*-value<br>if applicable |
| --- | --- | --- | --- |
|  | H | *p*-value |  |
| IL-6 | 1.77 | 0.67 | - |
| IL-8 | 1.97 | 0.63 | - |
| GM-CSF | 4.74 | 0.21 | - |
| MIP | 0.64 | 0.92 | - |

**Table S5:** Results from Wilcoxon signed rank test for Figure 6a

| Specification<br>including 0h | Wilcoxon signed rank test<br>hypothetical median = 1<br>*p*-value |
| --- | --- |
| *RUNX2* | *>0.99* |
| *SPP1* | *0.25* |
| *MMP2* | *0.50* |
| *PGK1* | *0.75* |
| *LDHA* | *0.50* |
| *VEGF* | *0.50* |
| *IL8* | *>0.99* |
| *IL6* | *0.50* |

**Table S6:**
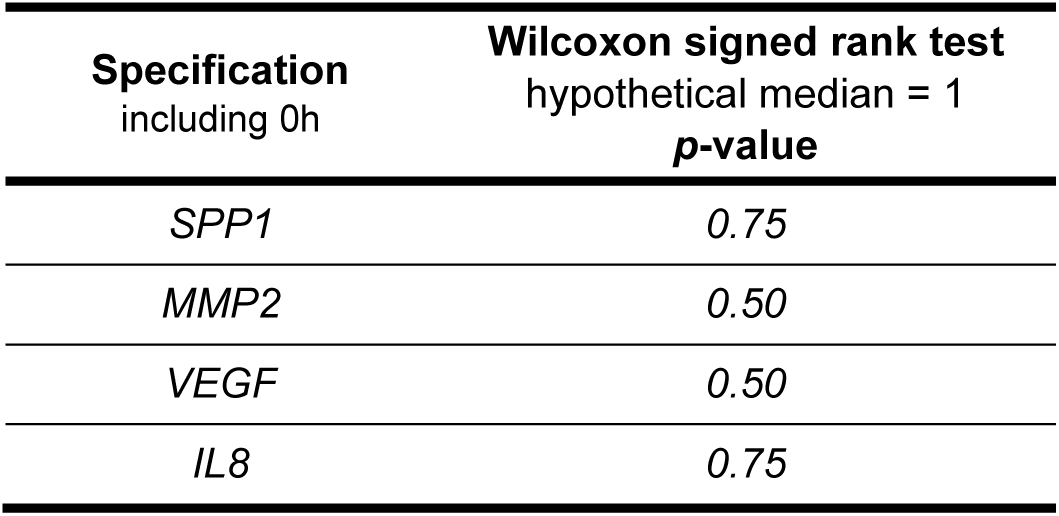
Results from Wilcoxon signed rank test for Figure 6b

**Table S7:** Results from Wilcoxon signed rank test for Figure 6c

| Specification<br>including 0h | Wilcoxon signed rank test<br>hypothetical median = 1 |
| --- | --- |
|  | *p*-value |
| *RUNX2* | 0.50 |
| *SPP1* | 0.50 |
| *MMP2* | 0.25 |
| *PGK1* | 0.75 |
| *LDHA* | >0.99 |
| *VEGF* | 0.75 |
| *IL8* | 0.25 |
| *IL6* | 0.25 |

**Table S8:** Results from Wilcoxon signed rank test for Figure 6d

| Specification<br>including 0h | Wilcoxon signed rank test<br>hypothetical median = 1 |
| --- | --- |
|  | *p*-value |
| *SPP1* | 0.25 |
| *MMP2* | >0.99 |
| *VEGF* | 0.75 |
| *IL8* | >0.99 |

**Table S9:** Results from Kruskal Wallis test with Dunn’s multiple comparisons test for Figure 6c

| Specification | Kruskal-Wallis test |  | Adjusted *p*-value<br>if applicable |
| --- | --- | --- | --- |
|  | H | *p*-value |  |
| IL-6 | 1.15 | 0.81 | - |
| IL-8 | 2.54 | 0.52 | - |
| GM-CSF | 2.69 | 0.49 | - |
| MIP | 1.68 | 0.68 | - |

**Table S10:**
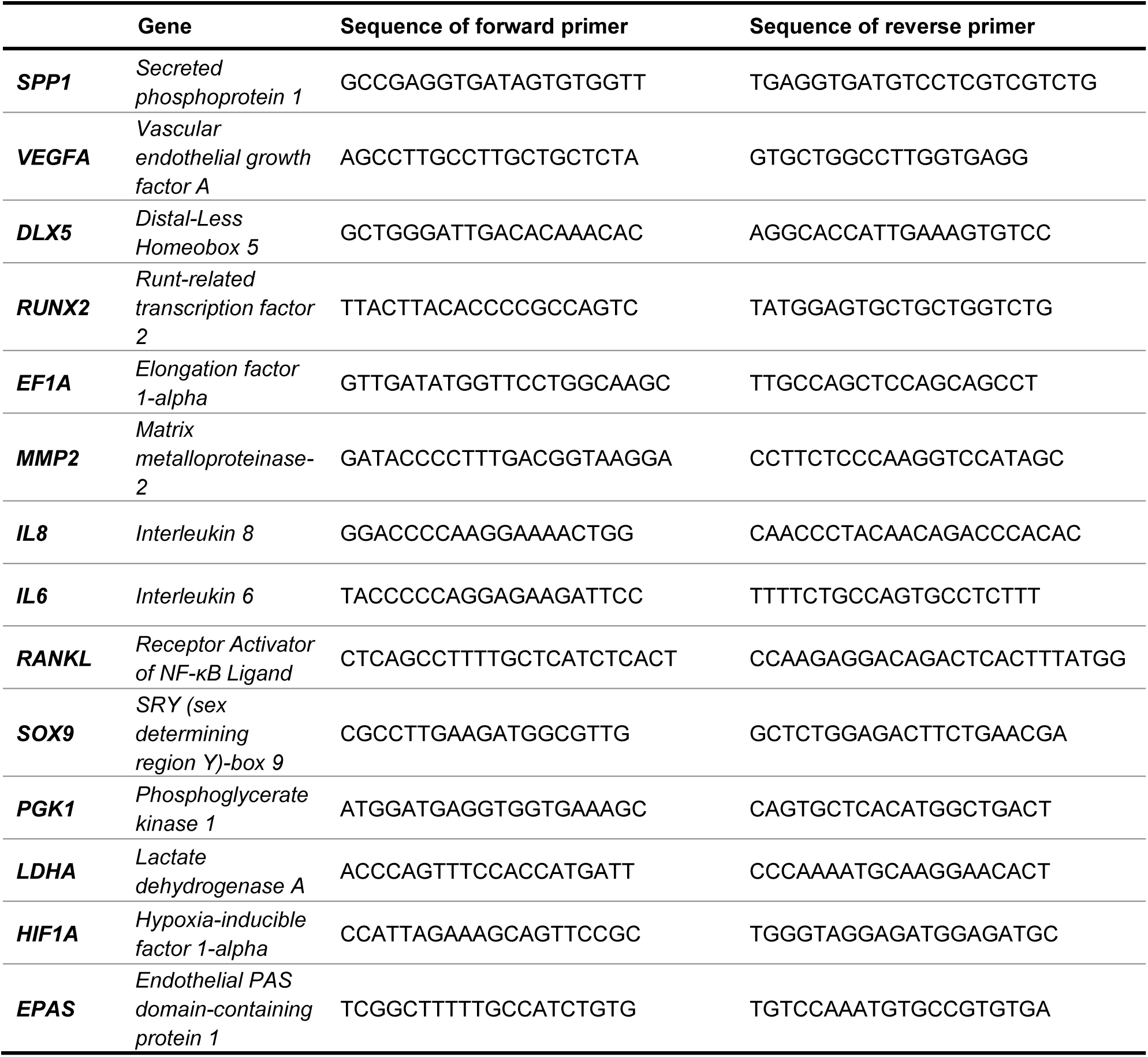
Sequences of primers used for qPCR.

